# *De novo* transcriptome assembly and the effect of foreign RNA contamination

**DOI:** 10.1101/2022.11.07.515468

**Authors:** Roberto Vera Alvarez, David Landsman

## Abstract

Multiple next-generation-sequencing (NGS)-based studies are enabled by the availability of a reference genome of the target organism. Unfortunately, several organisms remain unannotated due to the cost and complexity of generating a complete (or close to complete) reference genome. These unannotated organisms, however, can also be studied if a *de novo* reference transcriptome is assembled from whole transcriptome sequencing experiments. This technology is cost effective and widely used but is susceptible to off-target RNA contamination. In this manuscript, we present GTax, a taxonomy structured database of genomic sequences that can be used with BLAST to detect and remove foreign contamination in RNA sequencing samples before assembly. In addition, we investigate the effect of foreign RNA contamination on a *de novo* transcriptome assembly of *Solanum lycopersicum* (tomato). Our study demonstrates that removing foreign contamination in sequencing samples reduces the number of assembled chimeric transcripts.

## Introduction

Whole-genome and transcriptome sequencing has resulted in a greatly improved understanding of the biological complexities within organisms. Although whole-genome sequencing (WGS) is affordable for organisms with small genomes, it remains an expensive and complex task for organisms with larger genomes with more repetitive sequence regions.(1) Nevertheless, whole-transcriptome sequencing (WTS), also known as RNA sequencing (RNA-Seq), is a cost-effective means(2) to study differential gene expression profiles,(3, 4) phylogenomics,(5, 6) or plant evolution.(7, 8) It is particularly useful to create suitable reference transcriptomes for unannotated organisms using computational approaches called *de novo* transcriptome assemblies.(9) Assembled transcripts are annotated through the identification of homologous genes, proteins, and functional domains that could be cross-referenced with other public databases, such as Gene Ontology (GO).(10)

The lack of a reference genome of several species that have significant public health, economic, and environmental importance is a barrier in many studies. For example, green plants (*Viridiplantae* kingdom) or corals (*Anthozoa* class) are important groups of organisms but have limited annotations in public databases. The National Center for Biotechnology Information (NCBI) Genome database(11) contains 265 annotated reference genomes for the *Viridiplantae* kingdom but only 9 for the *Anthozoa* class. The NCBI Taxonomy database, however, shows 241,798 taxa in the *Viridiplantae* kingdom(12) and 8,252 taxa in the *Anthozoa* class(13) (as of February 22, 2022).

RNA-Seq contamination has played an important role in misleading multiple research conclusions (see the Background subsection in the Results and Discussion section below). Such contamination is more troublesome when the target organism does not have a reference genome with annotation available in public databases. Here, we present GTax, a taxonomic structured database of genomic sequences that can be used with BLAST for taxonomic classification and filtering. This approach efficiently detects and eliminates contaminant reads in RNA-Seq data. In addition, we discuss the effects of RNA-Seq contamination on *de novo* transcriptome assembly that uses the Trinity pipeline. We evaluate the quality of five *de novo* transcriptome assemblies for *Solanum lycopersicum* (tomato).

The first transcriptome was assembled using randomly selected reads from the tomato reference transcriptome. The second transcriptome was assembled by adding to the previous reads two million contaminant reads (20%) randomly selected from eight GTax taxonomy groups. Finally, the last three transcriptomes were assembled from 10 tomato wild-type RNA-Seq samples with different contamination levels. Assemblies were compared to the reference genome and transcriptome using Benchmarking Universal Single-Copy Orthologs (BUSCO) and rnaQuast.

## Results and Discussion

### Background

There are three main companies that provide RNA-Seq technologies: Illumina, Oxford Nanopore Technologies (ONT), and Pacific BioSciences (PacBio). Illumina remains the dominant RNA-Seq platform, as the company reports extremely low error rates and is affordable for high-sequence coverage depths.(14) Illumina-generated RNA-Seq short reads, however, can produce artifactual chimeras and fragmented transcripts during *de novo* transcriptome assembly.(1) ONT and PacBio sequencing technologies are designed to generate RNA-Seq long reads that could be used to sequence full-length transcripts. Nevertheless, these technologies result in high error rates and lower throughput. Alternative approaches have been developed using hybrid *de novo* transcriptome assembly, including both short and long RNA-Seq reads. These methods improve the quality of the assembly,(15) but the cost of using multiple sequencing platforms is a limitation to its general applicability.(16)

Multiple assemblers (e.g., Trinity,(17) Trans-ABySS,(18) SPAdes(19)) process Illumina RNA-Seq short reads. Trinity is the most commonly used tool for *de novo* transcriptome assembly. It was developed specifically for transcriptome assembly and uses de Bruijn graphs to generate multiple isoforms of a gene. In addition, Trinity offers an *in-silico* normalization method to process samples with different sequencing depths. Holzer and Marz compared assemblers and reported that these tools outperformed others; no tool, however, delivered perfect results for all analyzed data sets.^2^

As noted above, RNA-Seq short read-based assemblers generate transcriptomes with fragmented or chimeric isoforms. Therefore, additional steps are needed to identify spurious transcripts and assess the quality of assemblies. Multiple approaches have been developed to reduce false positive transcripts assembled *de nov*o.(2) When no reference genome is available for the target organism this ability remains problematic. Transcript abundance is quantified by calculating transcripts per million (TPM). The low-expression contigs with TPM levels that are lower than a cutoff value are discarded. NCBI BLAST tools(20) are used to identify spurious transcripts by sequence homology searches against multiple databases. BUSCO(21, 22) and rnaQuast(23) can be used to evaluate the quality of the assembly. BUSCO estimates the completeness and redundancy of an assembly based on universal single-copy orthologs. rnaQuast is intended for testing different assembly methods and pipelines on well-known organisms.

Theoretically, assemblers expect RNA-Seq data from a single organism. Therefore, the quality of a *de novo* assembly depends not only on the computational pipeline but also on the quality and purity of the RNA-Seq data. Contamination, however, is more common than expected in RNA-Seq samples;(24, 25) it is particularly problematic in samples from organisms for which there is no reference genome by which to frame the analysis.(26) Contamination of genomic and transcriptomic data can be classified as foreign and cognate. Foreign RNA-Seq contamination involve reads that originate from off-target, contaminant organisms, and cognate are reads that originate from off-target RNA species.(9)

Contamination has been the source of many inaccurate findings (e.g., a report of high rate of horizontal gene transfer (HGT) found in the tardigrade genome(27)). HGT was later rejected by Koutsovoulos et al. after finding bacterial contamination in the data.(28) Downstream analysis, such as the inference of phylogenomic trees, produce wrong classifications of taxonomies.(29) Inaccurate assemblies create bias in the analysis of a non-dietary origin of exogenous plant miRNAs reported to cross the mammalian gastrointestinal track.(30) Further, a common assumption is that contamination is *not* a problem when a reference genome exists(31); however, many studies have demonstrated that reference genomes should be used prudently due to existing contamination in public databases.(8, 25, 32, 33)

Detecting and removing contamination prior to WGS or WTS data analysis is a critical step in a pipeline for *de novo* transcriptome assembly. Ballenghien et al. stated, “Bioinformatic pipelines for NGS-based population genomic data should be further developed/improved in order to account for the probable existence of between-species and within-species contamination.”(26) We address this important issue in this paper.

Detecting the contamination in RNA-Seq short reads is complex due to sequence similarity between genes in distant taxonomic species. An illustrative example is the photosynthesis-related genes found in genomes of phototrophic bacteria but that originate from plants.(34) BLAST tools can be used to align RNA-Seq short reads to public databases of sequences to associate reads with one or more taxonomies. This is time and resource consuming, however, even when using modern cloud-computing platforms.(35) K-mer-based methods have been developed to accelerate the computation but at the cost of reducing the sensitivity of the taxonomy classification. Although k-mer-based tools are reported to be 900 times faster than BLAST, the latter is a more sensitive tool for sequence similarity identification.(36) Kraken2,(37) a k-mer-based tool, can be used for detecting contamination in RNA-Seq short reads but it depends greatly on the inclusion of the genome in the database(s). CLARK(38) is another tool that uses a supervised sequence classification with discriminative k-mers. CONSULT(39) tests whether k-mers from a query fall within a user-specified distance of the reference dataset using locality-sensitive hashing. Kaiju(40) executes a taxonomic classification for high-throughput sequencing reads but cannot be used to extract contaminant reads from raw sequence files. Conterminator(25) detects contamination in nucleotide and protein sequence sets using an all-against-all sequence comparison. Other tools for detecting RNA-Seq data contamination are FastQ Screen,(41) which uses Bowtie and BWA but is limited by the existence of a reference genome, and RNA-QC-Chain,(42) which can remove rRNA reads and identify foreign species in the sample using Hidden Markov Model searches but is incapable of identifying and removing foreign contaminant reads. For a detailed comparison among these tools, see Ounit et al.(38) and Cornet.(43)

Public databases at host institutes of the International Nucleotide Sequence Database Collaboration include a taxonomic classification for genome and transcriptome deposited data. The NCBI Sequence Read Archive(44) (SRA) uses an *in-house* developed taxonomic classification tool named Sequence Taxonomic Analysis Tool (STAT).(45) STAT is a scalable k-mer-based tool for fast assessment of taxonomic diversity intrinsic to SRA submissions. Although it offers valuable information and metadata, it was not designed for distribution. Downloading raw reads filtered by selected taxonomic identifiers from SRA archives is not possible.

### GTax, a taxonomic structured database of genomic sequences

The identification of contaminant reads in RNA-Seq samples is complex, especially when the source of contamination is unknown. The use of traditional sequence similarity search tools, such as BLAST, are inefficient in identifying contamination in raw data files, which may contain from 10 to 100 million reads. Public BLAST databases, such as NT and NR, have grown to more than 400GB of compressed indexes, making the screening of millions of short sequences impractical. As of February 22, 2022, the NT database contained 79,893,405 sequences and 674,286,748,964 total bases (UNIX command: blastdbcmd -info -db nt), and the NR database contained 460,326,648 sequences and 175,993,717,455 total residues (UNIX command: blastdbcmd -info -db nr). Because BLAST remains the most sensitive tool for sequence similarity searches, we designed a practical database for the taxonomic classification of short RNA-Seq reads.

NCBI released a new tool, Datasets,(46) that gathers data from across NCBI databases using command line instructions. This tool can be used to query the NCBI Genome databases and retrieve all available assemblies. It also provides machine-readable metadata that can be used for classification and filtering. We used Datasets metadata for the creation of GTax, a taxonomic structured database of genomic sequences that includes a subset of reference genomes, if available, or the latest assembly. Sequences were filtered by RefSeq Accession prefixes(47) to reduce the size and possibly contaminated sequences (see Methods section for details). The sequences were organized into 19 mutually exclusive and hierarchical taxonomic groups (**Error! Not a valid bookmark self-reference**.). For example, taxonomies in the *Viridiplantae* kingdom are divided into three GTax groups, *Liliopsida* includes all monocotyledon sequences, the Eudicotyledons group includes all dicotyledon sequences, and other taxa in the *Viridiplantae* kingdom not in these groups are placed in the *Viridiplantae* group. The same principle is applied to the *Chordata* phylum and all taxonomy groups from *Neoteleostei* to *Sarcopterygii*. Finally, all remaining *Eukaryote* taxa are placed in the Eukaryota taxonomy group.

This taxonomic structured division of the genomic sequences in GTax keeps phylogenetically closely related species in the same taxonomy group and significantly reduces the size of the searchable BLAST database. The *Sauropsida* group, which is the biggest group, contains 1,073 sequences and 46,172,754,879 total bases, only 6.84% of the NT database. The current version of GTax represents 72.18% of the sequences in the NT database.

Our taxonomic classification workflow assumes that RNA-Seq reads from a target organism aligned with the correct taxonomy group when the target organism has a reference genome or an assembly at any level included in GTax; these reads are identified as “correct” reads. If there are no sequences for the target organism in GTax, but genomic sequences from a phylogenetically closely related species are present, then some percent of the reads will align with the correct taxonomy group. Finally, if the target organism does not have genomic sequences nor phylogenetically closely related species in GTax, the “correct” reads will be classified as unidentified after a BLAST search against the rest of the GTax taxonomy groups.

Contaminant reads from organisms with genomic sequences in GTax align with their respective taxonomy groups. Those reads are identified as contamination and marked for removal. This approach identifies contaminant reads for known organisms with sequences available in GTax. Most of the common contaminants, such as bacteria, fungi, or human, can be identified using the GTax taxonomy groups. This approach, however, will not identify contaminant reads when information about the source is not available in public databases.

Our approach is initiated by screening (i.e., using BLAST searches) the RNA-Seq reads against the taxonomy group of the target organism. In these cases, we can screen millions of RNA-Seq reads against less than 6% of the NT database. Then, unidentified reads are screened against the rest of the GTax taxonomy groups. Reads labeled as “correct” are those that match the taxonomy group of the target organism. Those that remain unidentified are labeled as such.

We tested our approach in two ways. First, we selected 15 organisms from 12 taxonomy groups with reference genomes included in the GTax database. We generated overlapping single end reads, 100bp long, sequentially from the reference transcripts with an overlap window of 50bp. For the two *Bacteria* included, the reads were generated from the reference genome. Second, 15 RNA-Seq samples were selected from the SRA database for organisms without a reference genome, in addition to a highly contaminated human sample and a WGS sample from *Pseudomonas fluorescens*.

Tables 2A and 2B show the groups of “generated 100bp overlapped reads” from organisms with reference genomes. The correct taxonomy group for each organism is identified with a red background. The numbers in those cells are the percentages of the total reads generated from the reference transcriptome (genome for the two *Bacteria*); the numbers for the other, white background cells are the number of reads.

**Table 1:** GTax database content

| Taxonomy group | No. of taxonomies | Sequences | Size (GB) |
| --- | --- | --- | --- |
| Bacteria | 8,558 | 16,137 | 35.11 |
| Archaea | 336 | 554 | 0.90 |
| Liliopsida | 21 | 265 | 26.83 |
| Eudicotyledons | 79 | 880 | 39.17 |
| Viridiplantae | 12 | 184 | 4.52 |
| Fungi | 82 | 797 | 1.70 |
| Arthropoda | 101 | 1,364 | 26.06 |
| Neoteleostei | 75 | 1,437 | 44.60 |
| Actinopterygii | 37 | 1,047 | 45.52 |
| Glires | 25 | 2,178 | 30.44 |
| Primates | 20 | 433 | 39.32 |
| Carnivora | 31 | 286 | 28.89 |
| Artiodactyla | 28 | 447 | 36.90 |
| Amphibia | 9 | 122 | 30.36 |
| Sauropsida | 61 | 1,073 | 46.17 |
| Sarcopterygii | 29 | 229 | 26.83 |
| Chordata | 11 | 301 | 13.99 |
| Eukaryota | 71 | 803 | 8.76 |
| Viruses | 11,071 | 13,555 | 0.44 |

**Table 2A:** Alignment with each GTax taxonomic group of 100bp overlapped sequences generated from the reference transcriptome of several organisms

| Organism | Escherichia coli str. K-12<br>substr. MG1655 | Granulicella sp.<br>5B5 | Zea mays | Arabidopsis<br>thaliana | Solanum<br>lycopersicum | Pyricularia oryzae<br>70-15 | Zymoseptoria tritici<br>IPO323 |
| --- | --- | --- | --- | --- | --- | --- | --- |
| Assembly | GCF_000005845.2 | GCF_014083945.1 | GCF_902167145.1 | GCF_000001735.4 | GCF_000188115.4 | GCF_000002495.2 | GCF_000219625.1 |
| Taxonomy group in GTax | Bacteria | Bacteria | Liliopsida | Eudicotyledons | Eudicotyledons | Fungi | Fungi |
| Reads | 92,810 | 157,144 | 2,498,207 | 1,552,497 | 1,553,449 | 445,220 | 288,692 |
| Bacteria | 99.96 | 99.92 | 0 | 0 | 340 | 0 | 0 |
| Archaea | 0 | 0 | 0 | 0 | 0 | 0 | 0 |
| Liliopsida | 0 | 0 | 98.58 | 0 | 0 | 0 | 1 |
| Eudicotyledons | 0 | 0 | 7 | 98.51 | 98.53 | 0 | 0 |
| Viridiplantae | 0 | 0 | 4 | 0 | 0 | 0 | 0 |
| Fungi | 0 | 0 | 0 | 0 | 0 | 98.73 | 99.25 |
| Arthropoda | 0 | 0 | 1 | 0 | 0 | 0 | 0 |
| Neoteleostei | 0 | 0 | 0 | 0 | 0 | 0 | 0 |
| Actinopterygii | 0 | 0 | 0 | 0 | 0 | 0 | 0 |
| Glires | 0 | 0 | 0 | 0 | 0 | 0 | 0 |
| Primates | 0 | 0 | 0 | 0 | 0 | 0 | 0 |
| Carnivora | 0 | 0 | 0 | 0 | 0 | 0 | 0 |
| Artiodactyla | 0 | 0 | 0 | 0 | 0 | 0 | 0 |
| Amphibia | 0 | 0 | 0 | 0 | 0 | 0 | 0 |
| Sauropsida | 0 | 0 | 0 | 0 | 0 | 0 | 0 |
| Sarcopterygii | 0 | 0 | 0 | 0 | 0 | 0 | 0 |
| Chordata | 0 | 0 | 0 | 0 | 0 | 0 | 0 |
| Eukaryota | 0 | 0 | 0 | 0 | 0 | 0 | 0 |
| Viruses | 0 | 0 | 0 | 0 | 0 | 0 | 0 |

*Note.* Percentage of the reads aligned with the organism's taxonomic group are highlighted in red. Total number of reads in the "Reads" row.

**Table 2B:**
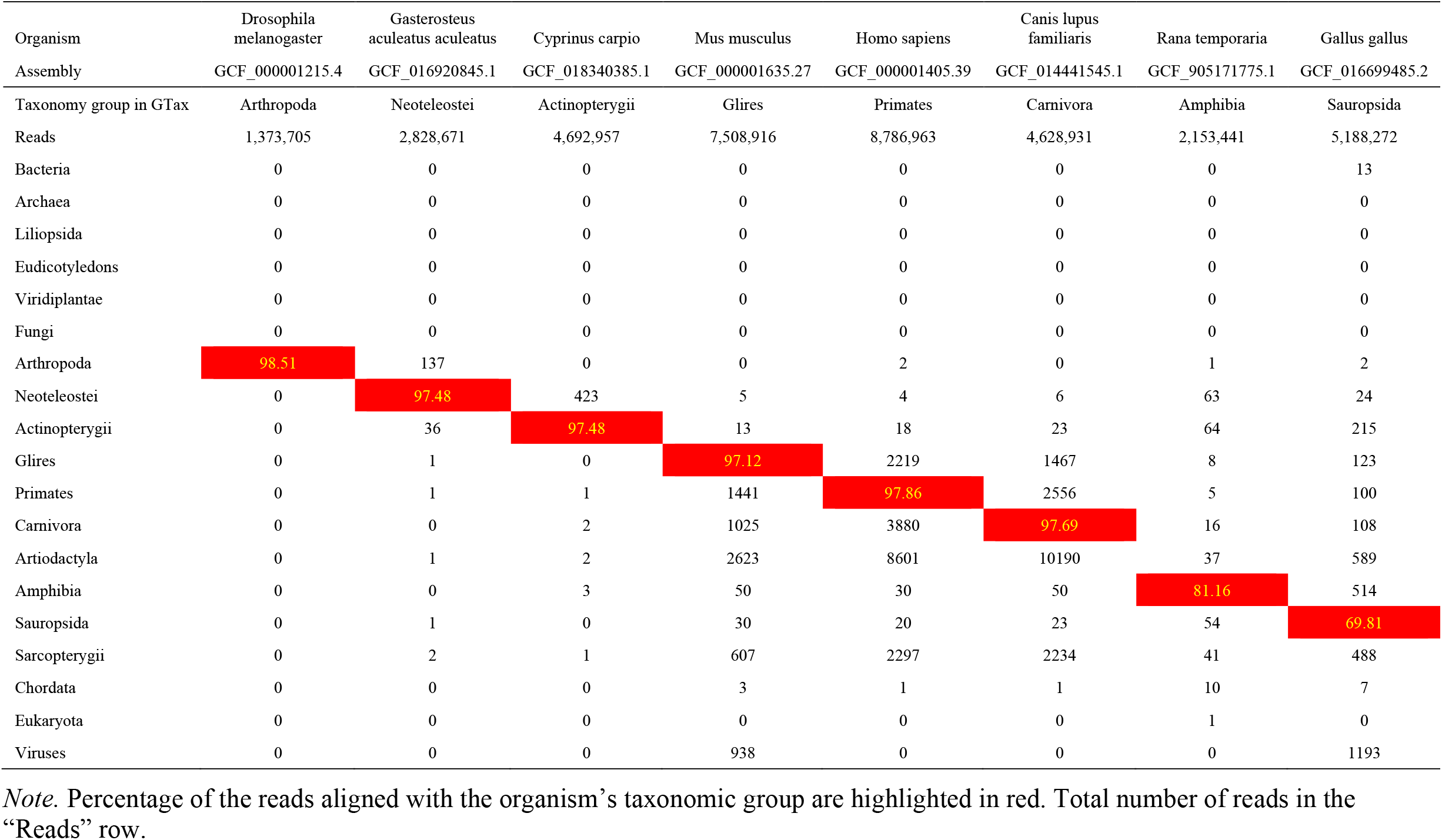
Alignment with each GTax taxonomic group of 100bp overlapped sequences generated from the reference transcriptome of each organism

| Organism | Drosophila melanogaster | Gasterosteus aculeatus aculeatus | Cyprinus carpio | Mus musculus | Homo sapiens | Canis lupus familiaris | Rana temporaria | Gallus gallus |
| --- | --- | --- | --- | --- | --- | --- | --- | --- |
| Assembly | GCF_000001215.4 | GCF_016920845.1 | GCF_018340385.1 | GCF_000001635.27 | GCF_000001405.39 | GCF_014441545.1 | GCF_905171775.1 | GCF_016699485.2 |
| Taxonomy group in GTax | Arthropoda | Neoteleostei | Actinopterygii | Glires | Primates | Carnivora | Amphibia | Sauropsida |
| Reads | 1,373,705 | 2,828,671 | 4,692,957 | 7,508,916 | 8,786,963 | 4,628,931 | 2,153,441 | 5,188,272 |
| Bacteria | 0 | 0 | 0 | 0 | 0 | 0 | 0 | 13 |
| Archaea | 0 | 0 | 0 | 0 | 0 | 0 | 0 | 0 |
| Liliopsida | 0 | 0 | 0 | 0 | 0 | 0 | 0 | 0 |
| Eudicotyledons | 0 | 0 | 0 | 0 | 0 | 0 | 0 | 0 |
| Viridiplantae | 0 | 0 | 0 | 0 | 0 | 0 | 0 | 0 |
| Fungi | 0 | 0 | 0 | 0 | 0 | 0 | 0 | 0 |
| Arthropoda | 98.51 | 137 | 0 | 0 | 2 | 0 | 1 | 2 |
| Neoteleostei | 0 | 97.48 | 423 | 5 | 4 | 6 | 63 | 24 |
| Actinopterygii | 0 | 36 | 97.48 | 13 | 18 | 23 | 64 | 215 |
| Glires | 0 | 1 | 0 | 97.12 | 2219 | 1467 | 8 | 123 |
| Primates | 0 | 1 | 1 | 1441 | 97.86 | 2556 | 5 | 100 |
| Carnivora | 0 | 0 | 2 | 1025 | 3880 | 97.69 | 16 | 108 |
| Artiodactyla | 0 | 1 | 2 | 2623 | 8601 | 10190 | 37 | 589 |
| Amphibia | 0 | 0 | 3 | 50 | 30 | 50 | 81.16 | 514 |
| Sauropsida | 0 | 1 | 0 | 30 | 20 | 23 | 54 | 69.81 |
| Sarcopterygii | 0 | 2 | 1 | 607 | 2297 | 2234 | 41 | 488 |
| Chordata | 0 | 0 | 0 | 3 | 1 | 1 | 10 | 7 |
| Eukaryota | 0 | 0 | 0 | 0 | 0 | 0 | 1 | 0 |
| Viruses | 0 | 0 | 0 | 938 | 0 | 0 | 0 | 1193 |

*Note.* Percentage of the reads aligned with the organism's taxonomic group are highlighted in red. Total number of reads in the "Reads" row.

Table 2A includes the first group: *Bacteria, Green Plants*, and *Fungi*. More than 98% of the reads from these organisms are aligned with the correct taxonomy group. In the specific case of *Solanum lycopersicum* (tomato), 340 reads are aligned with the Bacteria taxonomy group, which may indicate some level of contamination in the transcriptome.

The second group of organisms, presented in Table 2B, is different. Although more than 97% of the generated reads align with the correct taxonomy group for all organisms except *Rana temporaria* (frog) and *Gallus gallus* (chicken), there is an increased number of reads aligned with other taxonomy groups, indicating a varied amount of contamination. The frog and chicken examples show a lower number of reads aligned with the correct taxonomy group but also few reads aligned with other groups. We suspect that, in addition to some contamination, these transcriptomes include some chimeric transcripts that are not aligned with any genomic sequence. Chimeric transcripts also are the most probable explanation for the small percentage of generated reads that remain unidentified in all examples. After further investigation, we confirmed that all unidentified reads belong to computationally predicted transcripts (accessions prefixes with XR_ and XM_) included in the transcriptomes and are probably not valid (for more details on RefSeq prefixes, see NCBI RefSeq Accession prefixes(47)).

These experiments demonstrate that “correct” reads can be identified in high numbers when the target organism has a reference genome, or an assembly, included in GTax. Tables 3A and 3B provide the results for the circumstances when the target organism sequences are not included in GTax. These tables include a row in yellow with the percentage of filtered reads that can be used to assemble the transcriptome after removal of contaminant sequences. Cells colored with the same background, the correct taxonomy group, and the remaining unidentified reads for each organism (columns) are summed to generate “Percentage of reads for assembly” row (yellow in all organisms).

**Table 3A:** Samples from SRA database for organisms without a reference genome

| Organism | *Pseudomonas fluorescens* | *Cylindrospermopsis raciborskii* FACHB-1096 | *Lolium perenne* | *Physalis peruviana* | *Opuntia streptacantha* | *Diplocarpon rosae* | *Cimex lectularius* | *Synodus* sp. isolate FZ12 FC-2018 |
| --- | --- | --- | --- | --- | --- | --- | --- | --- |
| Sample | SRR5823570 | SRR16571653 | SRR3340606 | SRR1952996 | SRR3478177 | SRR5178307 | SRR3297746 | SRR8242436 |
| Taxonomy group in GTax | Bacteria | Bacteria | Liliopsida | Eudicotyledons | Eudicotyledons | Fungi | Arthropoda | Neoteleostei |
| Reads | 2,302,977 | 12,431,662 | 11,062,381 | 21,845,419 | 41,904,616 | 658,128 | 8,527,345 | 6,811,855 |
| Reads for assembly | 99.99 | 99.15 | 99.26 | 98.74 | 95.90 | 99.39 | 97.95 | 99.28 |
| Unidentified | 7.64 | 62.90 | 93.59 | 62.34 | 92.97 | 59.86 | 93.69 | 95.22 |
| Bacteria | 92.35 | 36.25 | 0.02 | 0.35 | 3.77 | 0.13 | 0.02 | 0.00 |
| Archaea | 0.00 | 0.00 | 0.00 | 0.00 | 0.00 | 0.00 | 0.00 | 0.00 |
| Liliopsida | 0.00 | 0.01 | 5.67 | 0.04 | 0.13 | 0.01 | 0.01 | 0.00 |
| Eudicotyledons | 0.00 | 0.05 | 0.01 | 36.40 | 2.93 | 0.40 | 0.03 | 0.00 |
| Viridiplantae | 0.00 | 0.00 | 0.00 | 0.02 | 0.06 | 0.00 | 0.00 | 0.00 |
| Fungi | 0.00 | 0.54 | 0.00 | 0.80 | 0.00 | 39.53 | 0.01 | 0.00 |
| Arthropoda | 0.00 | 0.00 | 0.02 | 0.04 | 0.00 | 0.05 | 4.26 | 0.03 |
| Neoteleostei | 0.00 | 0.02 | 0.01 | 0.00 | 0.00 | 0.00 | 0.05 | 4.06 |
| Actinopterygii | 0.00 | 0.00 | 0.02 | 0.00 | 0.00 | 0.00 | 0.02 | 0.46 |
| Glires | 0.00 | 0.02 | 0.04 | 0.00 | 0.02 | 0.00 | 1.79 | 0.02 |
| Primates | 0.00 | 0.17 | 0.33 | 0.00 | 0.10 | 0.00 | 0.02 | 0.01 |
| Carnivora | 0.00 | 0.00 | 0.00 | 0.00 | 0.00 | 0.00 | 0.01 | 0.01 |
| Artiodactyla | 0.00 | 0.02 | 0.00 | 0.00 | 0.00 | 0.00 | 0.02 | 0.01 |
| Amphibia | 0.00 | 0.00 | 0.00 | 0.00 | 0.00 | 0.00 | 0.01 | 0.01 |
| Sauropsida | 0.00 | 0.00 | 0.00 | 0.00 | 0.00 | 0.00 | 0.01 | 0.06 |
| Sarcopterygii | 0.00 | 0.00 | 0.00 | 0.00 | 0.00 | 0.00 | 0.03 | 0.09 |
| Chordata | 0.00 | 0.00 | 0.00 | 0.00 | 0.00 | 0.00 | 0.00 | 0.00 |
| Eukaryota | 0.00 | 0.00 | 0.00 | 0.01 | 0.00 | 0.01 | 0.03 | 0.00 |
| Viruses | 0.00 | 0.00 | 0.28 | 0.00 | 0.00 | 0.00 | 0.00 | 0.00 |

*Note.* Values expressed in percentages. Reads in cell of the same color for each sample are summed to generate the final reads to assembly.

**Table 3B:** Samples from SRA database for organisms without a reference genome

| Organism | Acipenser sp | Cavia porcellus | Homo sapiens | Ursus americanus | Eptesicus fuscus | Spea bombifrons | Taeniopygia guttata | Influenza A virus |
| --- | --- | --- | --- | --- | --- | --- | --- | --- |
| Sample | SRR16661141 | SRR12442784 | SRR16958449 | SRR14160197 | SRR4249968 | SRR9160217 | DRR185733 | SRR7734450 |
| Taxonomy group in GTax | Actinopterygii | Glires | Primates | Carnivora | Artiodactyla | Amphibia | Sauropsida | Viruses |
| Reads | 21,742,680 | 24,000,000 | 16,818,866 | 17,092,718 | 25,227,832 | 21,336,768 | 17,778,851 | 9,731,528 |
| Reads to assembly | 99.73 | 99.67 | 22.17 | 99.02 | 93.00 | 93.35 | 99.97 | 22.94 |
| Unidentified | 24.70 | 5.18 | 2.14 | 25.66 | 85.61 | 86.43 | 10.39 | 12.24 |
| Bacteria | 0.04 | 0.00 | 77.78 | 0.00 | 0.00 | 2.28 | 0.00 | 0.09 |
| Archaea | 0.00 | 0.00 | 0.00 | 0.00 | 0.00 | 0.00 | 0.00 | 0.00 |
| Liliopsida | 0.01 | 0.00 | 0.00 | 0.01 | 0.01 | 0.50 | 0.00 | 0.03 |
| Eudicotyledons | 0.00 | 0.00 | 0.00 | 0.00 | 0.01 | 1.96 | 0.00 | 0.26 |
| Viridiplantae | 0.00 | 0.00 | 0.00 | 0.00 | 0.00 | 0.00 | 0.00 | 0.01 |
| Fungi | 0.00 | 0.00 | 0.00 | 0.00 | 0.00 | 0.03 | 0.00 | 0.00 |
| Arthropoda | 0.00 | 0.00 | 0.00 | 0.03 | 0.04 | 0.13 | 0.00 | 0.10 |
| Neoteleostei | 0.07 | 0.00 | 0.00 | 0.01 | 0.07 | 0.48 | 0.01 | 2.27 |
| Actinopterygii | 75.02 | 0.02 | 0.00 | 0.01 | 0.06 | 0.46 | 0.00 | 0.44 |
| Glires | 0.04 | 94.50 | 0.03 | 0.23 | 1.13 | 0.42 | 0.00 | 5.48 |
| Primates | 0.01 | 0.17 | 20.03 | 0.34 | 1.61 | 0.12 | 0.00 | 22.48 |
| Carnivora | 0.01 | 0.05 | 0.01 | 73.36 | 1.81 | 0.03 | 0.00 | 45.75 |
| Artiodactyla | 0.00 | 0.04 | 0.01 | 0.23 | 7.40 | 0.03 | 0.00 | 0.02 |
| Amphibia | 0.06 | 0.00 | 0.00 | 0.00 | 0.01 | 6.92 | 0.00 | 0.10 |
| Sauropsida | 0.02 | 0.00 | 0.00 | 0.01 | 0.06 | 0.17 | 89.57 | 0.00 |
| Sarcopterygii | 0.00 | 0.03 | 0.00 | 0.11 | 2.16 | 0.04 | 0.00 | 0.02 |
| Chordata | 0.01 | 0.00 | 0.00 | 0.00 | 0.00 | 0.02 | 0.00 | 0.00 |
| Eukaryota | 0.00 | 0.00 | 0.00 | 0.00 | 0.00 | 0.01 | 0.00 | 0.00 |
| Viruses | 0.00 | 0.00 | 0.00 | 0.00 | 0.00 | 0.00 | 0.00 | 10.71 |

*Note.* Values expressed in percentages. Reads in cell of the same color for each sample are summed to generate the final reads to assembly.

Table 3A shows low levels of contamination for the RNA-Seq samples. For *Pseudomonas fluorescens* (SRR5823570), which has phylogenetically closely related species in GTax, 92.35% of the reads aligned with the correct taxonomy group, with 7.64% unidentified reads (these reads sum to 99.99% of the reads, are labeled as “correct,” and are used in the assembly step). In addition, 36.25% of reads from the other bacterium, *Cylindrospermopsis raciborskii FACHB-1096* (SRR16571653), are aligned with the correct taxonomy group, while 62.90% remain unidentified. Both *Fungi* and *Primates* sequences contaminate this sample. Overall, 99.15% of the reads can be used for assembly. Most of the reads for the three plant examples are unidentified, with some contamination detected from *Bacteria, Fungi*, and *Primates*. The Eudicotyledons examples, *Physalis peruviana* and *Opuntia streptacantha*, display a different level of reads aligned with the correct taxonomy group. For *Physalis peruviana*, which is closely related to tomato, 36.40% of the reads align with the Eudicotyledons taxonomy group, whereas *Opuntia streptacantha* does not have a closely related organism in the database, and most of the reads are unidentified (92.97%).

Table 3B shows the second group of analyzed samples. Similarly, organisms such as *Synodus sp. isolate FZ12 FC-2018* with a low number of reads aligned with the correct taxonomy group (Table 3A), and *Cavia porcellus*, closely related to mouse, has 94.50% of reads aligned with the correct taxonomy group (Table 3B).

The human sample included in this example (SRR16958449) contains 20.03% of the reads aligned with *Primates* and 2.14% of reads unidentified, for a total of 22.17% of the reads ready for assembly. This sample, however, contains a high level of bacterial contamination, 77.78% of the reads. (The SRA STAT report also similarly shows 78.45% of contaminated reads (see the Analysis tab in https://trace.ncbi.nlm.nih.gov/Traces/sra/?run=SRR16958449). This sample was extracted from vascular aortic smooth muscle cells, and no bacterial presence was reported in the study. All other samples in this study show similar bacterial contamination. The general approach includes the unaligned reads for further analysis. In cases with well-annotated reference genomes, such as human, mouse, yeast, or some bacteria, however, we recommend using reads aligned only with the target taxonomy group.

The *Influenza A virus* sample included in Table 3B displays a high number of reads aligned with the *Primates* and *Carnivora* taxonomy groups. Virus samples are usually collected from host organisms. In this case, *Mustela putorius furo* (domestic ferret), which belongs to the *Carnivora* taxonomy group, 22.48% of the reads in this samples align with *Primates*, indicating additional human contamination that is significant.

We have released a python package that can be used to generate the GTax taxonomy group FASTA files for the creation of BLAST indexes. Please see https://gtax.readthedocs.io/.

### Effect of RNA-Seq contamination on de novo transcriptome assembly

We used the *Solanum lycopersicum* (tomato) reference transcriptome and a wild-type tomato RNA-Seq dataset to study the effect of foreign contamination on *de novo* transcriptome assembly. The tomato genome is well annotated with a reference genome and transcriptome available (assembly ID GCF_000188115.4). The current annotated genome includes 45,901 transcripts (see https://www.ncbi.nlm.nih.gov/genome/?term=txid4081[orgn]).

Five tomato transcriptomes were assembled *de novo* using different pre-processing approaches. The first transcriptome assembly contains randomly extracted 100bp long reads from the tomato reference transcriptome. This was repeated four times to produce four paired-end samples of ∼9 million reads each. The second transcriptome was assembled with the same tomato reads as the first plus two million (∼20%) randomly selected contaminant reads added to each sample (4% *Bacteria*, 1% *Archaea*, 10% *Fungi*, 1% *Arthropoda*, 1% *Chordata*, 1% *Metazoan*, 1% *Eukaryote*, and 1% *Viruses*). We added more *Fungi* and *Bacteria*, as they are the most probable plant sample contaminants. It is important to note that these two sets of generated samples contain the same tomato reads. The only difference is the contaminant reads added to the second set.

The three other transcriptomes were assembled from ten tomato wild-type RNA-Seq samples selected from the SRA database (Supplementary File 2, tab: “Table 1 – Tomato WT samples”). The samples belong to four different BioProjects from different plant tissues. The first RNA-Seq transcriptome was assembled after trimming the adapters and filtering out low-quality reads (Trimmed assembly in Figure 1). The second RNA-Seq transcriptome was assembled with the trimmed reads that match the Eudicotyledons taxonomy group in GTax (*Eudicotyledons* assembly in Figure 1). The third RNA-Seq transcriptome was assembled with the Eudicotyledons-matched reads above plus the unidentified reads that remain after screening the samples against all other GTax taxonomy groups (Eudicotyledons + unidentified assembly in Figure 1).

**Figure 1:**
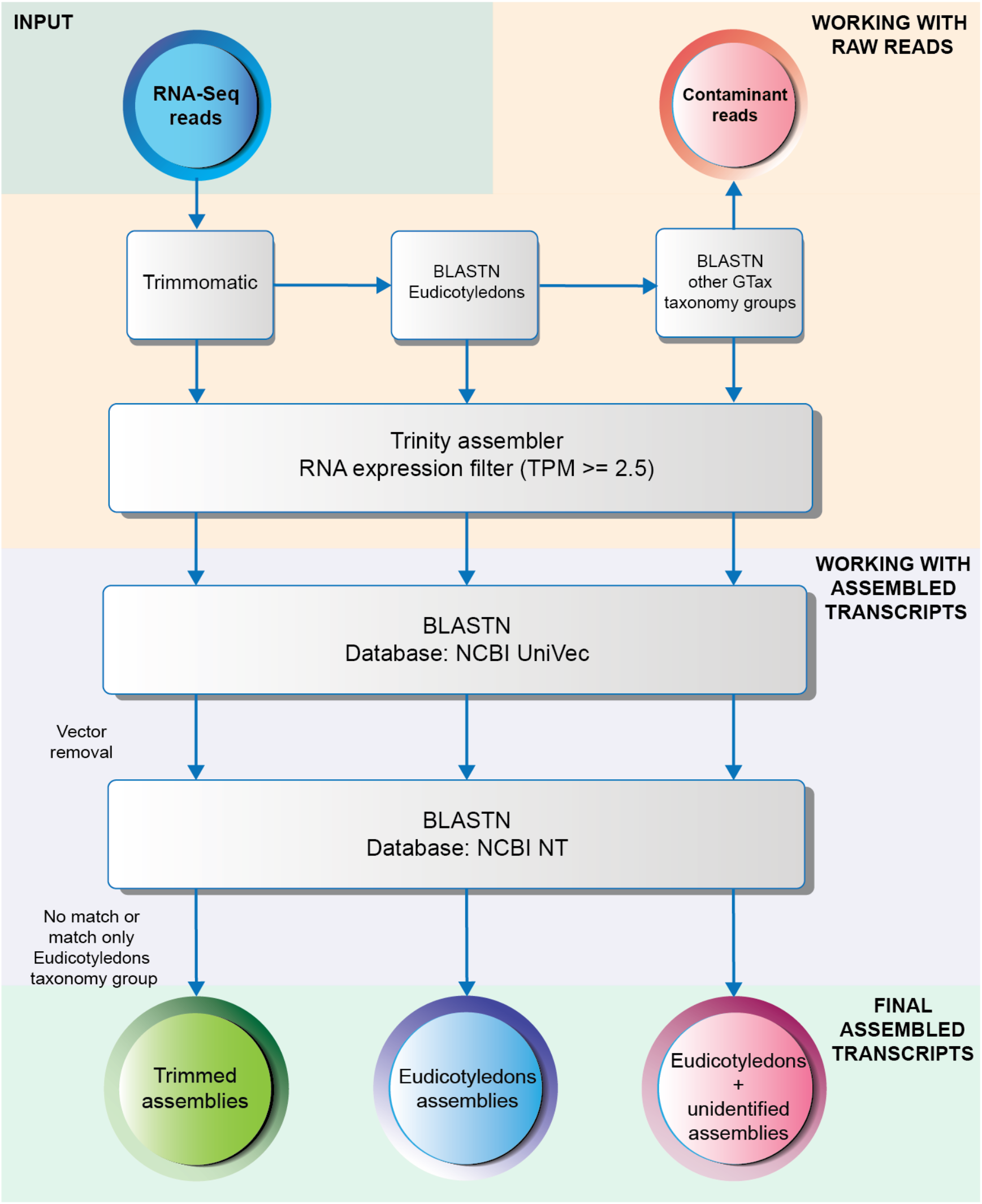
Workflow to remove vectors and contaminated transcripts after assembly completion. Different levels of decontamination of the SRA samples were used to assemble three transcriptomes: Trimmed, Eudicotyledons, and Eudicotyledons + unidentified.

After assembly, transcriptomes were filtered to remove lowly expressed transcripts using a TPM cutoff of ≥2.5. Two different post-assembly decontamination screenings were performed. First, transcripts were screened with BLASTN against the NCBI UniVec database(48) to detect and remove vectors. Then, a second BLASTN screen is done against the NT database to identify and remove contaminated transcripts (**Error! Reference source not found**.). This final BLAST screen is the starting point of the annotation process.(35)

To evaluate the quality of RNA-Seq assemblies, we used rnaQuast software.(23) Figure *2* shows the rnaQuast results of aligned transcripts for the five assemblies. The transcriptome assembled with the 100bp generated reads from the tomato reference transcriptome contains 39,232 transcripts aligned with the reference genome (38,610 uniquely and 22 multiply aligned). Although this assembly used reads generated from only the tomato reference transcriptome, only 16 contaminant transcripts were identified, using BLASTN against the NT database, and removed after assembly. As expected, these 16 transcripts aligned with *Bacteria* sequences (in GTax), supporting the assumption that bacterial contamination is minimal in the tomato reference transcriptome, as identified in Table 2A. Only three transcripts in the transcriptome were unaligned with the reference genome after all the post-processing steps. Transcripts identified in Figure *2* as unaligned are considered false positive transcripts (probably chimeric) and remain in the transcriptome. Thus, Trinity generates only three chimeric transcripts when using reads generated from only the tomato reference transcriptome.

**Figure 2:**
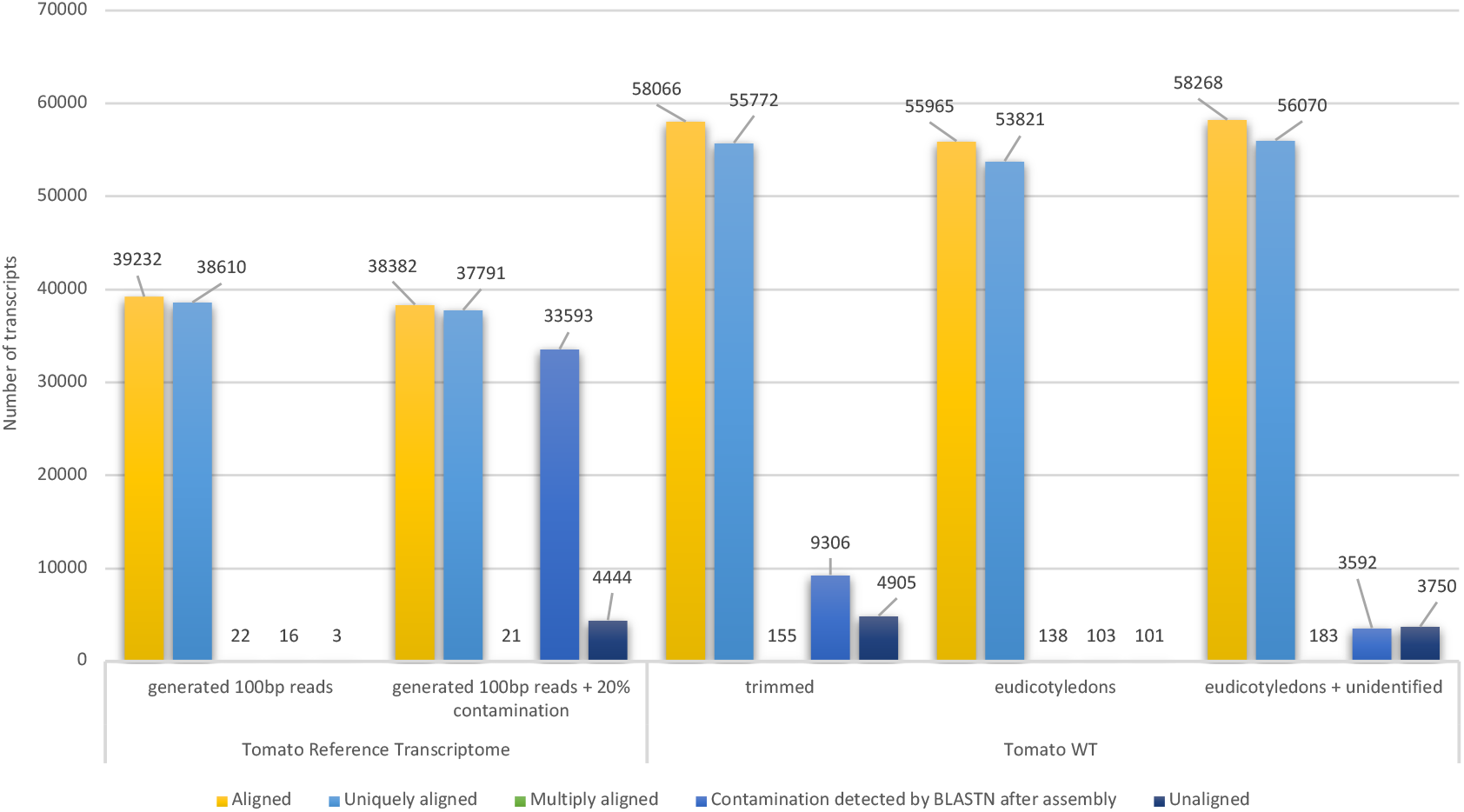
Alignment results reported by rnaQuast for five tomato transcriptomes *de novo* assembled in this study. (Supporting data for Figure 2 is in Supplemetary file 2).

The second transcriptome assembled shows a different distribution of transcripts. This transcriptome was assembled with the same tomato reads plus 20% contaminant reads from several other species. In this case, 850 fewer assembled transcripts align with the reference genome. Post-assembly decontamination steps remove 33,593 contaminated transcripts, but 4,444 unaligned transcripts remain in the transcriptome. The Trinity software seems to mix some reads from the contaminant sequences with tomato sequences to create chimeric transcripts absent in the previous assembly.

The last three transcriptomes using tomato wild-type RNA-Seq data corroborates the assumption that contamination affects the final assembly significantly. The transcripts assembled from the trimmed reads contain 4,905 unaligned transcripts that are reduced to 101 when only Eudicotyledons reads are used. This number increases to 3,750 when the unidentified reads are added to the samples. The Eudicotyledons assembly, however, includes 2,101 fewer transcripts aligned with the reference genome than does the trimmed assembly and 2,249 fewer than the Eudicotyledons + unidentified assembly.

rnaQuast reports an increase in the duplication ratio from 1.2 in the first two assemblies to 1.7 in the three RNA-seq samples, producing a difference of >16,000 transcripts (see Supplementary File 2, tab: “Table 2 – rnaQuast short report”). There seems to be a proportional relationship between duplicated transcripts and the number of reads used in the assembly.

We also assessed the exon coverage of the alignment of each assembled transcript set to the tomato genome. BLASTN was used to align the assembled transcripts to the tomato reference genome. BLAST high-scoring segment pairs (HSPs) were used to quantify the sequence coverage of the annotated transcripts by counting the overlap between HSPs and annotated exons. Figure 3 shows the number of transcripts that match annotated isoforms in the tomato reference genome. In all cases, there is a reduction in the number of transcripts that match annotated isoforms when contamination is present. Using the generated 100bp tomato reads, Trinity assembled 68.96% of annotated transcripts with >80% sequence coverage. This is reduced to 44.94% using the Eudicotyledons reads, which is the best coverage obtained with the SRA samples.

**Figure 3:**
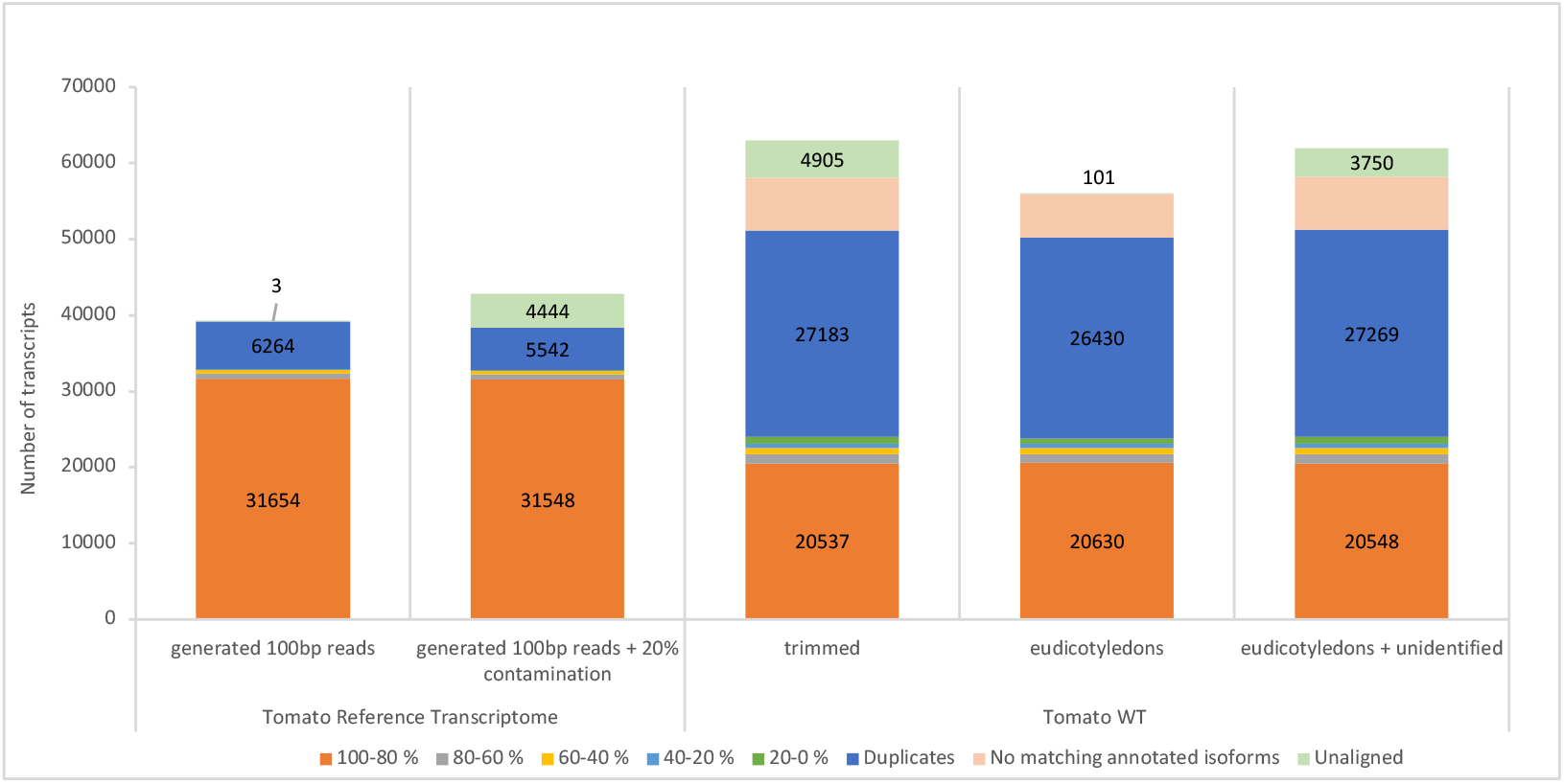
Percentage of sequence coverage of annotated isoforms is reported in color for each assembly. (Supporting data for Figure 3 is in Supplemetary file 2).

For the three RNA-Seq samples, fewer transcripts match isoforms with <80% coverage, showing a high completeness of the assembly. The SRA-based assemblies show, as reported by rnaQuast, high duplication levels. In addition, in these cases, >5,000 transcripts do not overlap any annotated transcript (Figure 3, category “No matching annotated isoform”). These transcripts are peculiar in that they align with the genome in a single BLAST HSP that covers the entire transcript. Most also show an expression level, or TPM value, close to the cutoff used to discard lowly expressed transcripts.

Figure 4 shows an example in a genomic context, where one of the unannotated transcripts (TRINITY_DN1213_c2_g1_i2) does not match any annotated isoform, and is aligned with an intron of TRINITY_DN876_c1_g1_i2 (in purple) in the same genomic region. The first one matched exactly to the annotated gene, LOC101262544. It is clearly validated by the RNA-Seq alignment coverage and the spanning reads (dark grey gaps between exons) in all SRA samples in this study. Samples SRR13931770 and SRR14575350 collected from the anther and fruit tissues, respectively, however, show some intronic sequence coverage that was used by Trinity to assemble TRINITY_DN1213_c2_g1_i2. We should clarify that these two SRA samples belong to different BioProjects and were collected independently. It is difficult to determine whether this is DNA contamination or an artifact of the experimental assembly protocol. In our opinion, this is a false positive transcript that should be eliminated from the final assembly. It is difficult or practically impossible, however, to detect this class of false positives when the target organism does not have a reference genome.

**Figure 4:**
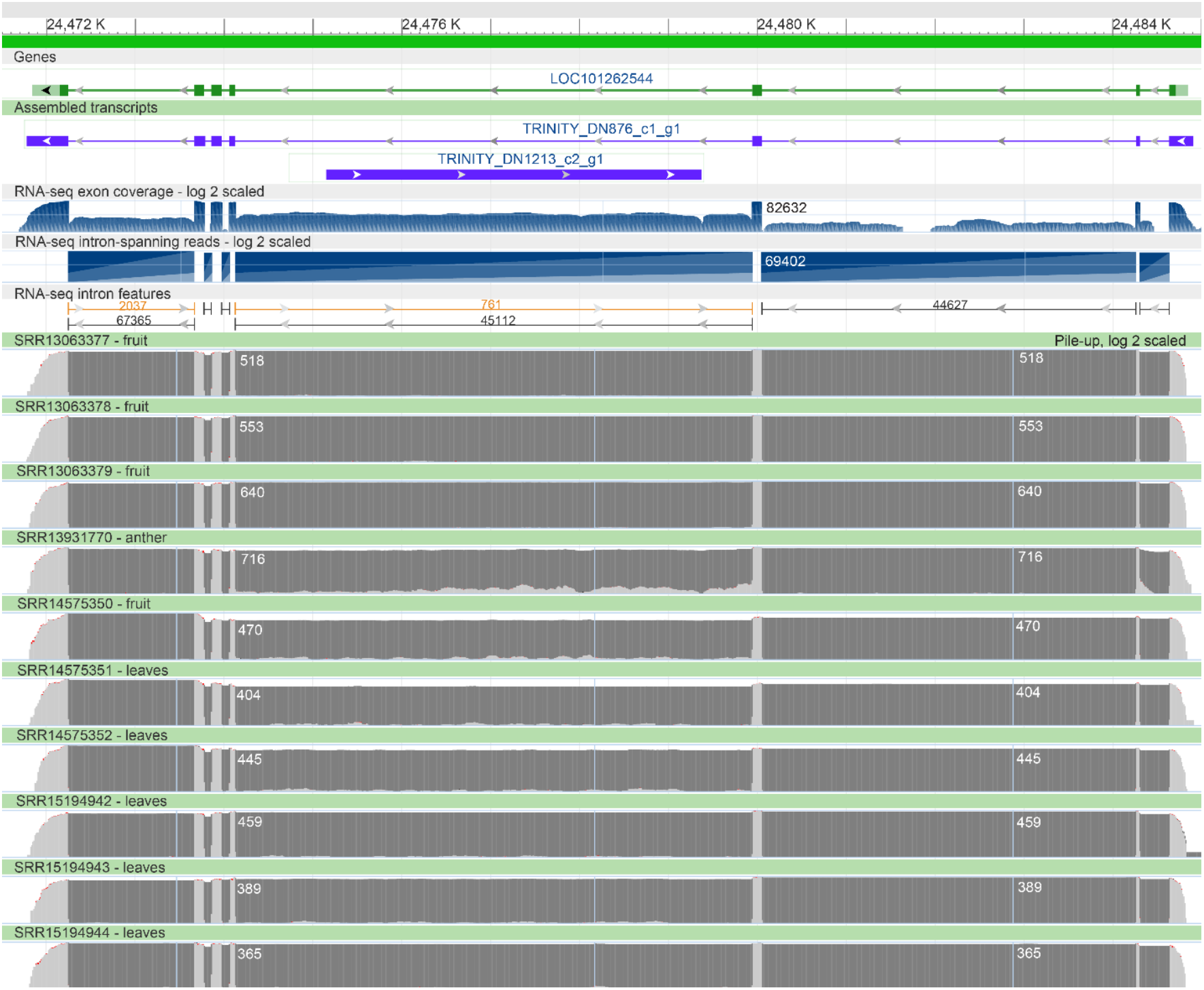
Genomic view for the annotated gene LOC101262544 (green) and aligned Trinity transcripts to the same genomic region: TRINITY_DN876_c1_g1_i2 and TRINITY_DN1213_c2_g1_i2 (purple). *Note:* Annotated exon coverage, intron spanning reads, and intro features; pile-up of alignment coverage in log_2_ scale for the SRA samples used in this study.

**Figure 5:**
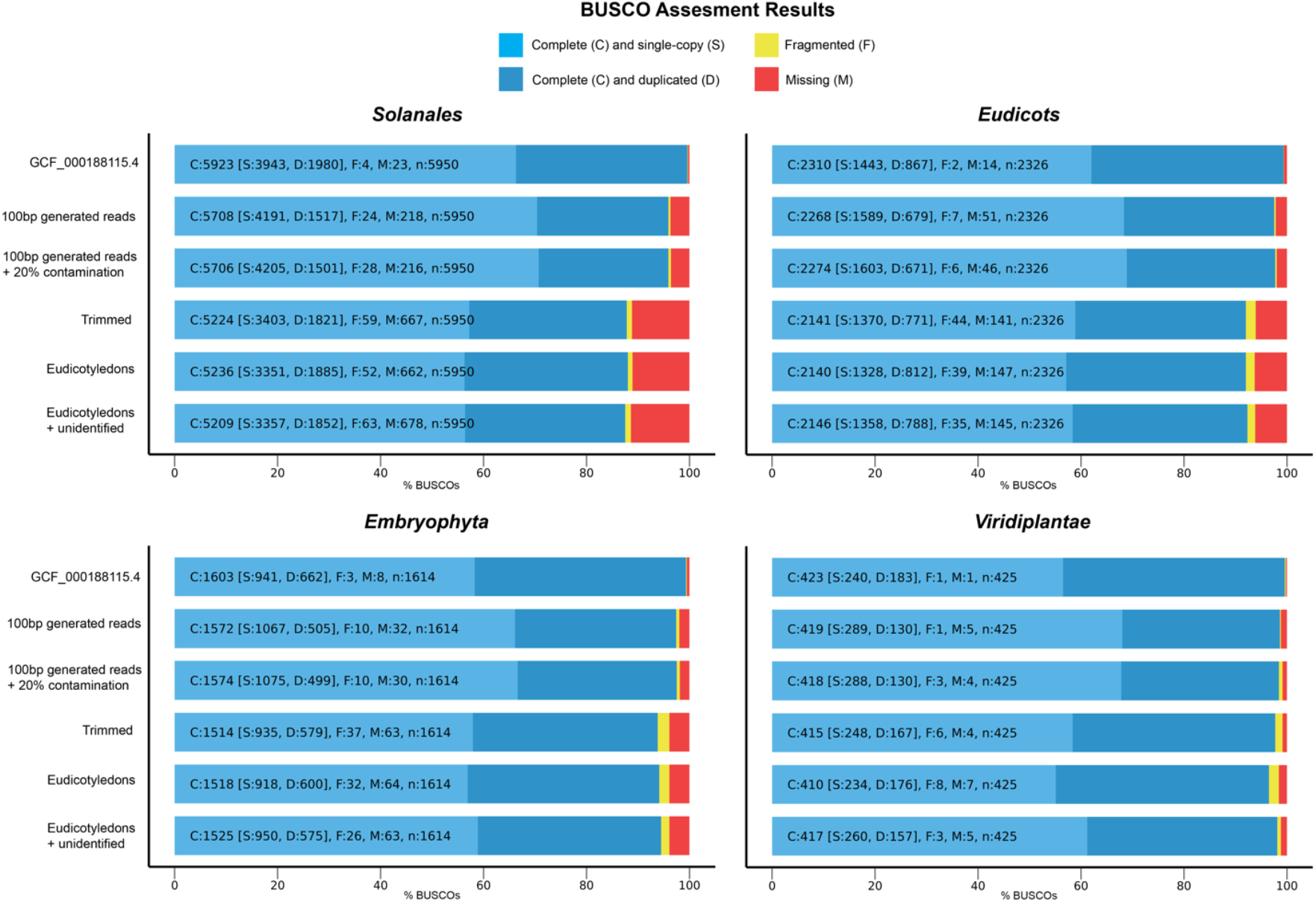
BUSCO profiles for the five assembled transcriptomes generated at different taxonomic levels.

We also performed BUSCO analyses to assess the completeness of the Trinity assemblies. BUSCO profile plots were generated for four taxonomic levels to compare the reference transcriptome and the five assemblies (**Error! Reference source not found**.). The profile plot created for the *Solanales* order shows that the 100bp generated assemblies are similar, with a difference of only two missing BUSCO profiles. The same differences are present when comparing the assemblies generated from the SRA samples. Although the Eudicotyledons assembly is less fragmented and is missing some BUSCO profiles, there is no significant difference in comparison to the other assemblies. The same pattern is present in the BUSCO plots for the *Eudicots* and *Embryophyta* clades. The Eudicotyledons assembly, however, is more fragmented and is missing BUSCO profiles in the *Viridiplantae* kingdom compared with the other two SRA-based assemblies.

The BUSCO plots demonstrate that RNA-Seq contamination does not affect the highly conserved genes at different taxonomic levels. Further, our decontamination steps, which remove contaminant and chimeric transcripts, do not affect the BUSCO completeness of the assembly.

After annotation of the assembled transcripts, the transcriptome assembly process is completed. Assembled transcripts can be annotated and cross-referenced with public databases, such as GO,(10) NCBI Conserved Domain Database,(49) COG,(50) Pfam,(51) UniProt,(52) and eggNOT.(53) Trinotate,(54) for instance, is a popular transcriptome assembly and annotation framework that uses Trinity for the assembly and most of the aforementioned databases for the annotation. This reduces the possibility of reporting chimeric transcripts as relevant biological entities. We also recommend prioritizing annotated transcripts when using *de novo* transcriptomes as a reference in differential gene expression analyses.

## Conclusions

WTS is a valuable technology to study a wide range of biological processes even if the target organism does not have a reference genome. In this case, *de novo* transcriptome assemblers, such as Trinity, can be used to produce reference transcriptomes with a high level of assembly completeness and specificity. These tools, however, generate some fragmented and chimeric transcripts that are difficult to identify without a reference genome. In our opinion, an assembly is completed when the transcripts are annotated and cross-referenced to public databases. These annotations can be used as extra validations of assembled transcripts. Foreign and cognate RNA-Seq contamination removal is a critical step in the assembly process. Although it is not included in most popular assembly pipelines, RNA-Seq contamination, if not removed, increases the number of chimeric transcripts, which affects downstream analysis. We recommend the use of GTax, a taxonomic structured database of genomic sequences, for detecting foreign contamination of transcriptome and genome sequencing data. Although we tested GTax with BLASTN, the database also can be searched with k-mer-based tools to speed up computation. A freely available python package can be used to generate a GTax database from the NCBI Genome database https://gtax.readthedocs.io/.

Transcriptome assembly is a complex process that requires the integration of many bioinformatics tools and methodologies usually in a pipeline. The assembler is not the only critical step; pre-processing steps to prepare the data and post-processing steps, such as vector detection, contamination removal, and final annotation, make a *de novo* assembly a viable transcriptome reference for further analysis.

## Methods

### GTax

Assembly metadata for four taxonomy superkingdoms (*Archaea, Bacteria, Viruses*, and *Eukaryotes*) were gathered using NCBI Datasets, version 12.19.0. Only RefSeq genomic sequences were used because, of the three main genomic data host institutes, NCBI is the only one that uses a contamination screening pipeline for WGS data submissions. Each superkingdom set of metadata was processed with an *in-house* developed and freely available python package (https://gtax.readthedocs.io/). The first step of the filtering process is to select, for each taxonomy, the reference genome, if available, or the latest assembly. Then, unplaced sequences inside the assemblies are discarded because most include contamination. Finally, sequence accessions starting with RefSeq prefixes such as **NW** and **NZ** were excluded, except for the case of **NZ_CM**, and **NZ_CP**, which are the codes for complete chromosomes in GenBank. GTax taxonomy groups were created with three files: FASTA, text file with the relationship between sequence accession and TaxID (used to create the BLAST databases with taxonomy information), and a final file with the same relationship plus the file offset where the sequence can be extracted directly.

### RNA-Seq processing

We used BLAST version 2.13.0+ to identify matches between the reads and GTax sequences. BLAST parameters used to define a match were (a) percentage of identity larger than 75%, (b) query (read) coverage larger than 75%, (c) e-value smaller than 1.0×10^−5^, and (d) the penalty for nucleotide mismatch equals -3. FASTQ files were transformed to FASTA and divided into files that contained 50,000 sequences each to speed up processing.

### Assemblies

Trinity version 2.13.2 with default parameters were used to generate the assemblies. Transcript quantification was executed as described in the manual (http://trinityrnaseq.github.io/). A TPM cutoff of 2.5 was used to filter out lowly expressed transcripts. BUSCO version 4.1.2 with databases odb10 were used to generate the BUSCO profiles, using default parameters. RNAQuast version 2.2.1 with default parameters were used to compare the assemblies generated in this study.

## Ethics approval and consent to participate

Not applicable.

## Consent for publication

Not applicable.

## Availability of data and materials

GTax is implemented as a Python package under Public Domain license. Source code available at: https://github.com/ncbi/gtax. The documentation is available at: https://gtax.readthedocs.io/

Current version of GTax FASTA files are available for download at: https://console.cloud.google.com/storage/browser/gtax-database

## Authors’ contributions

RVA and DL contributed to the design of all the experiments and the manuscript preparation. All authors read and approved the final manuscript.

## Acknowledgements

This work utilized the computational resources of the NIH HPC Biowulf cluster (http://hpc.nih.gov).

We thank the NCBI BLAST Group: Christiam Camacho, Vadim Zalunin, Greg Boratyn, Ryan Connor, and Tom Madden, for their use of BLAST.

